# Actin modulates shape and mechanics of tubular membranes

**DOI:** 10.1101/712505

**Authors:** A. Allard, M. Bouzid, T. Betz, C. Simon, M. Abou-Ghali, J. Lemière, F. Valentino, J. Manzi, F. Brochard-Wyart, K. Guevorkian, J. Plastino, M. Lenz, C. Campillo, C. Sykes

## Abstract

The actin cytoskeleton shapes cells and also organizes internal membranous compartments. In particular, it interacts with membranes in intracellular transport of material in mammalian cells, yeast or plant cells. Tubular membrane intermediates, pulled along microtubule tracks, are involved during these processes, and destabilize into vesicles. While the role of actin in this destabilization process is still debated, literature also provide examples of membranous structures stabilization by actin. To directly address this apparent contradiction, we mimic the geometry of tubular intermediates with preformed membrane tubes. The growth of an actin sleeve at the tube surface is monitored spatio-temporally. Depending on network cohesiveness, actin is able to stabilize, or maintain membrane tubes under pulling. Indeed, on a single tube, thicker portions correlate with the presence of actin. Such structures relax over several minutes, and may provide enough time and curvature geometries for other proteins to act on tube stability.

---

Membranes in cells are the boundaries of numerous internal compartments such as organelles and vesicles. Such membranous structures constantly reorganize during intracellular trafficking, a process ensuring the targeted movement of substances in the cell interior. For example, flat membrane surfaces within the endoplasmic reticulum mature into highly curved cylinders and spheres (*1*). In particular, trafficking involves tubular intermediates pulled by molecular motors walking on microtubules (*2*). These dynamical rearrangements of membranes motivated the identification and the study of specialized proteins that bind to the membrane and directly act on its curvature (*3–5*). However, the orchestration of morphological changes needs pauses and shape stabilization processes that are often neglected (*1*).

Moreover, experiments on living cells show that the actin cytoskeleton is associated with intermediate tubular membranes. But how actin is involved in tube fate remains an open question (*6, 7*). An attractive idea is that actin might play a role by applying physical forces and stresses, that could ultimately lead to tube scission. Alternatively, an actin layer could have a stabilizing effect on a membrane (*8*). These hypothesis are difficult to address in the complex intracellular environment. Besides the size of transport vesicles and width of membrane tubes, between ten to hundred nanometers, are comparable with the actin network meshsize. This questions whether actin polymerization could affect tube morphology.

Here, we isolate the role of the actin cytoskeleton on membrane tube morphology in a biomimetic assay made of a membrane tube at the surface of which we polymerize an actin network. The preformed membrane tube is extruded from a liposome and held by an optically trapped bead. Such tubes are stable under a non-zero point force 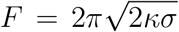, where *κ* is the membrane bending energy and *σ* the membrane tension (*9*). An adapted microinjection system sequentially delivers the activator of actin polymerization targeted to the membrane, then actin monomers that polymerize on the tube surface. The actin network is branched through the Arp2/3 complex, thus mimicking the situation in cells (*10, 11*). We show here that in the presence of an actin sleeve, a membrane tube can be stable even when the external pulling force vanishes. The membrane tube, surrounded by its actin sleeve, can be further pulled by the optical tweezers, at a force and a speed mimicking the pulling of membrane tubes by molecular motors walking on microtubules (*12*).

The actin network, made of entangled actin branches, produces a variety of outcomes under pulling, which depend on the thickness of the actin sleeve around the membrane tube. At a sleeve thickness higher than a few hundreds of nanometers, the network is unable to disentangle and a sheath of actin remains around the membrane tube that maintains its radius. We show that less that one minute of network growth is enough to obtain a stabilization of the tubular structure of the membrane that is robust and lasts for tens of minutes. At smaller sleeve thicknesses, discontinuous regions appear and smaller tube radii are observed in portions where the actin sleeve is absent. Therefore we never observe actin-induced scission of these membrane tubes, but rather, actin provides a way of modulating the radius of tubes along their length.

## Results

### Actin network growth around a membrane tube

We polymerize a branched actin network at the surface of a preformed membrane tube containing nickel lipids. A histidine-tagged version of pVCA, the proline rich domain-verprolin homology-central-acidic sequence from human WASP that activates the Arp2/3 complex (supplementary materials), binds to nickel lipids and activates actin polymerization at the tube surface. To avoid unnecessary actin polymerization in the solution we provide actin monomers only once the membrane is activated for polymerization. The time at which actin polymerization starts on the membrane tube is rigorously controlled by proceeding sequentially as follows. First, the preformed fluorescent membrane tube, maintained by an optically trapped bead, is bathed in a solution containing the other necessary proteins (P-solution with the Arp2/3 complex, profilin and capping protein, supplementary materials and Fig. 1A(a)). Second, pVCA is targeted to the membrane by microinjection close to the tube; proper injection is monitored by a sulforhodamine dye (Fig. 1A(b)). Third, actin monomers are microinjected and this moment sets the time (*t*_*i*_ in Fig. 1A(c)) at which actin polymerization starts around the tube. Actin becomes visible by fluorescence on the membrane after a few seconds (Fig. 1A(c)) and continues to grow (Fig. S1A). After 2 minutes, a sleeve of actin is obtained around the membrane tube (Fig. 1A(d)), whereas it fails to form when pVCA is omitted (Fig. S1, compare B and C). We define this composite system of membrane tube sheathed with an actin sleeve as “membrane-and-actin-sleeve” (MaAS).

**Fig. 1.**
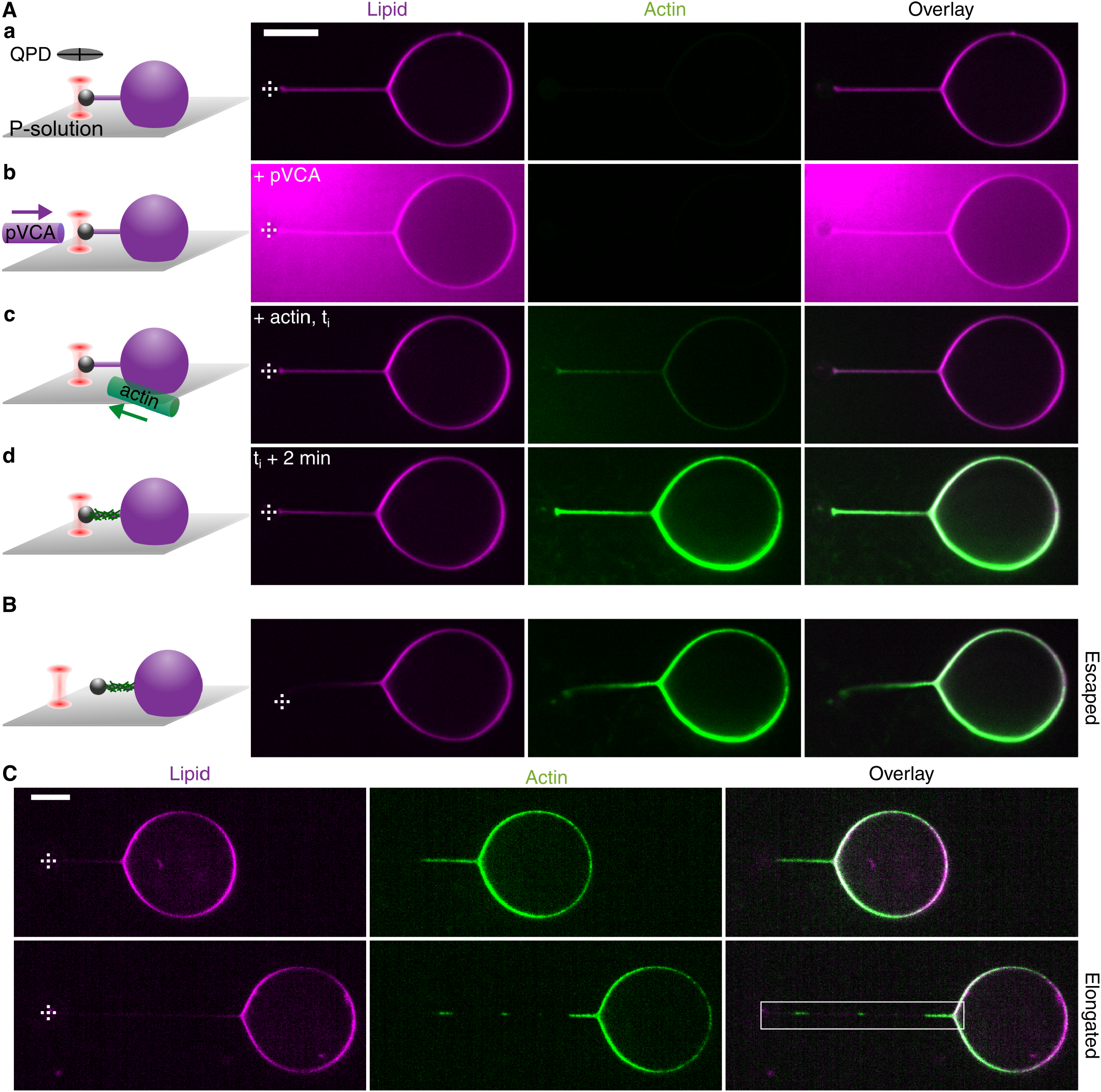
Effect of an actin sleeve on membrane tube stability. Lipids (magenta) and actin (green) are observed by spinning disk confocal imaging. (A and B) Left column: scheme of each step towards MaAS formation. (A) (a) Preformed tube held by optical tweezers, (b) microinjection of pVCA in a sulforhodamine-B solution, (c) microinjection of monomeric actin at *t*_*i*_, (d) the membrane tube is sheathed with an actin sleeve within 2 minutes. (B) Escaped MaAS and (C) elongated MaAS before (top) and after (bottom) pulling; the white box indicates the location of the elongated MaAS. Scale bars: 10 µm. Dashed crosses indicate bead center.

### MaAS characteristics under pulling forces comparable to the cellular situation

To mimic tubular intermediates pulled by molecular motors walking on microtubules, we subject MaAS to elongation by moving the stage at a controlled speed of 0.5 *−* 4 µm/s to reach tube lengths of 15 *−* 30 µm. We observe two cases. The first is an escape of the bead from the optical trap after a short MaAS elongation of less than 1 µm. In this case, the MaAS, thereafter called “escaped MaAS”, retains its shape indicating that the rigidity of the actin sleeve is strong enough to hold the bead in position even outside the trap (Fig. 1B). The tube does not retract, and this lasts for tens of minutes (Fig. S2). What we observe strikingly differs from a membrane tube in the absence of actin that would totally retract and reincorporate in the liposome within hundreds of milliseconds (*13*). In the second case, a MaAS continuously elongates during stage displacement, and is thereafter called “elongated MaAS” once the stage is stopped (Fig. 1C). Note that the actin sleeve in elongated MaAS might be discontinuous. We then explore whether the amount of actin in the sleeve may be different in these two cases.

### Actin thickness imposes MaAS fate

We visualize lipids and actin by confocal microscopy and concomitantly record the force *F* in the limit of 50 pN on the optically trapped bead (supplementary materials). First, we quantify the amount of actin by measuring the total actin fluorescence intensity of MaAS before pulling, normalized by the membrane tube fluorescence (supplementary materials). The actin content is clearly higher in escaped MaAS than in elongated MaAS (Fig. 2A). Second, force-elongation curves of escaped MaAS reveal a linear dependence of the force *F* as a function of the tube elongation *∆ℓ* (filled circles, Fig. 2B, supplementary materials). Escaped MaAS therefore appear linearly elastic (*F* = *k∆ℓ*), with an average spring constant *k* = 31 *±* 6 pN/µm. A linear force-extension curve is also observed for elongated MaAS and naked tubes with drastically lower slopes, respectively 0.38 *±* 0.11 pN/µm and 0.30 *±* 0.09 pN/µm (Fig. 2B).

**Fig. 2.**
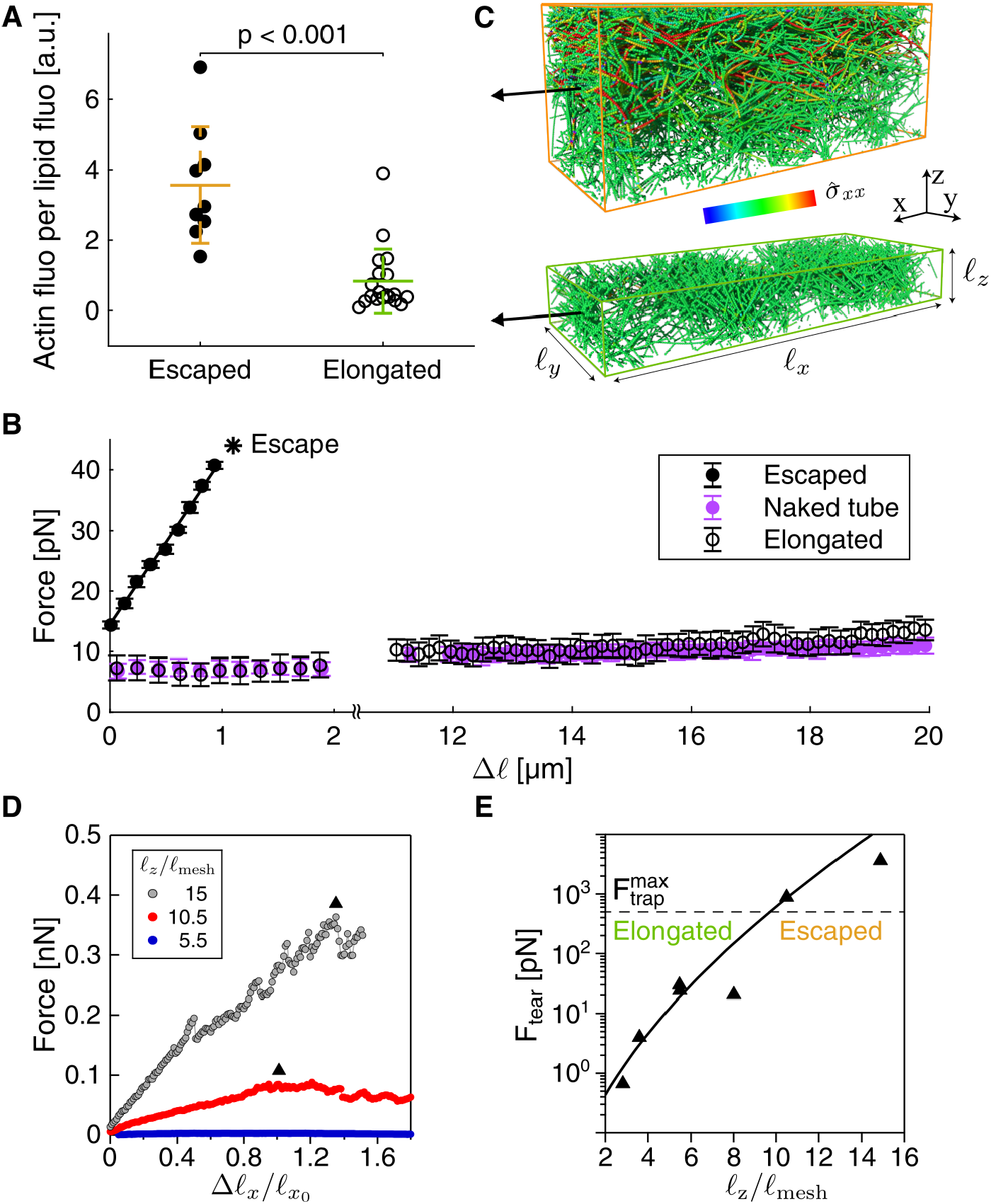
Actin sleeve thickness drive MaAS fate. (A, B) Escaped (filled circles, n = 8) and elongated (opened circles, n = 17) MaAS. Data shown as mean + s.d.. (A) Quantification of actin fluorescence per lipid fluorescence depending on MaAS fate. p-values calculated using the t-test. (B) Force-elongation curves for MaAS and a naked tube (light magenta filled circles, n = 15). The star symbol indicates the length at which the MaAS escapes. (C) Snapshots of two branched actin network networks for different thicknesses *ℓ*_*z*_ (top *ℓ*_*z*_ = 10.5*ℓ*_mesh_, bottom *ℓ*_*z*_ = 5.5*ℓ*_mesh_) under uniaxial deformation along the *x* axis. The two configurations correspond to a deformation of *∼* 150%. Colors indicate the local longitudinal stress 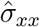 at the scale of a monomer, ranging from red for tension to blue for compression. (D) Average force *F* extrapolated for a cylindrical sleeve with inner radius *R*_0_ = 25 nm and variable thickness *ℓ*_*z*_, implying an outer radius *R*_0_ + *ℓ*_*z*_ as a function of the deformation *∆ℓ*_*x*_/*ℓ*_*x0*_ = (*ℓ*_*x*_ − *ℓ*_*x0*_)/*ℓ*_*x0*_, with *ℓ*_*x0*_ the initial size, for different network thicknesses. Triangles indicate the maximum force *F*_tear_ that the network can bear before falling apart. (E) Phase diagram representing *F*_tear_ as a function of the gel thickness. The dashed black line shows the optically trapped bead force limit of about 50 pN. Actin sleeves larger than *∼* 10 meshsizes should thus display an escaped MaAS behavior.

Escaped MaAS are robust elastic structures where the membrane tube remains under the sleeve. The membrane tube is held by the bead that is 3.05 µm in diameter, much larger than the membrane tube diameter. Therefore, full tube retraction is retained by the bead and the presence of the actin sleeve. In these conditions, escaped MaAS behavior is dominated by the contribution of the actin network. Therefore the thickness of the actin sleeve can be estimated from the elastic spring constant *k*. The actin network elastic modulus *E* was measured previously in similar experimental conditions as 10^3^ *−* 10^4^ Pa (*14*). The spring constant *k* of the actin sleeve is related to the sleeve cross-section area *S*, MaAS length *L*, and *E* as *k* = *ES/L*. With *L* = 15 *−* 30 µm and our experimental measurement of *k* ≃ 31 pN/µm, we find *S* = 5 *×* 10^*−*14^ *−* 9 *×* 10^*−*13^ m^2^ that leads to a sleeve radius of 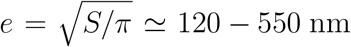. Note that we neglect here the hollowness of the sleeve which has a contribution to the section area that is one order of magnitude lower for a typical radius of 25 nm in our experiments.

### Simulation of a branched actin network under deformation

To account for the difference in MaAS fate depending on the actin content of the actin sleeve (Fig. 2A), we perform detailed molecular dynamics simulation of a entangled branched actin network under deformation (supplementary materials). Physically, our experimental observations suggest that thin actin sleeves are less cohesive than their thicker counterparts, and thus tend to fall apart to yield an elongated MaAS. Conversely, we reason that actin filaments in thicker sleeves are more extensively entangled, implying a more robust structure resulting in an escaped MaAS. To validate this picture and determine the minimal actin thickness required for an escaped MaAS, we apply a uniaxial quasi-static deformation along the *x* axis of simulated networks with different thicknesses (Fig. 2C). While internal local stresses remain low in thin networks, large and strongly heterogeneous stresses develop in thick networks, since entangled filaments pull on one another to maintain network cohesion (Fig. 2C, compare bottom to top). Assuming an actin persistence length of 10 µm (*15*) and a network meshsize *ℓ*_mesh_ = 30 nm (*16*) in the same protein mix as here, we derive the corresponding force-extension curves for a whole cylindrical actin sleeve of inner radius 25 nm and variable thickness *ℓ*_*z*_, which is shown in Fig. 2D. While all networks display an initial linear response, in thin networks, the force peaks at a relatively modest value *F*_tear_, following which the network looses its cohesion and the force decreases. By contrast, thick networks display much larger tearing forces *F*_tear_. Fig. 2E compares this tearing force to our maximum tweezing force of 50 pN (dashed line) for different values of network thicknesses *ℓ*_*z*_. For any sleeves whose tearing force exceeds the maximum tweezing force, the optical tweezers will give in before the sleeve does, and based on Fig. 2E we thus predict that any sleeve thicker than *∼* 10 meshes results in an escaped MaAS, while a thinner sleeve yields an elongated MaAS. This corresponds to a critical sleeve radius of *e ∼* 300 nm, consistent with our experimental estimate above.

Interestingly, this phase diagram reveals that the rigidity of the actin network increases almost exponentially as a function of the number of meshes. Our findings highlight that a small variation of the amount of actin, through a difference in a few units of meshes, displaces the system efficiently from a stable to an unstable state.

### Morphology of elongated MaAS

A striking observation in elongated MaAS is that the membrane tube is continuously present whereas the actin sleeve appears discontinuous (Fig. 3A). Interestingly, higher actin fluorescence along the tube correlates with higher lipid fluorescence, revealing that a thicker membrane tube is present under stable actin sleeve regions (Fig. 3A). To quantify this effect, for each MaAS, we define the membrane tube radius *r*_M_ where the actin signal is maximal, and *r*_m_ where the actin signal is minimal (including equals to zero) along the tube (respectively “M” and “m” in Fig. 3A and Fig. S3A-C). With *r*_0_ the membrane tube radius before the MaAS is pulled, the relative variation of radius reads *δr/r*_0_ = (*r*_M_ *− r*_m_)*/r*_0_ and is obtained directly from lipid fluorescence intensities (Fig. 3A and Fig. S3A-C and supplementary materials). Moreover, elongating a MaAS reveals three different situations that are sketched in Fig. 3B: in two of them, the sleeve maintains roughly its initial length but sits either next to the bead (Fig. 3A and Fig. 3B-*a*) or next to the liposome (Fig. 3B-*b*); in the third situation, the actin sleeve extends together with the membrane tube (Fig. 3B-*c*) The heterogeneity along the tube, through the ratio *δr/r*_0_, is close to zero before pulling (“Ref” condition, Fig. 3B) and increases when the MaAS is elongated, revealing the presence of radius heterogeneity along the tube (*a*, *b* and *c*, Fig. 3B). According to our classification, this increase is significant for the *b*-type, where the actin sheath stays close to the liposome. Independently of the (*a*, *b* and *c*) classification, the relative variation of radius increases in a high actin content situation (Fig. 3C).

**Fig. 3.**
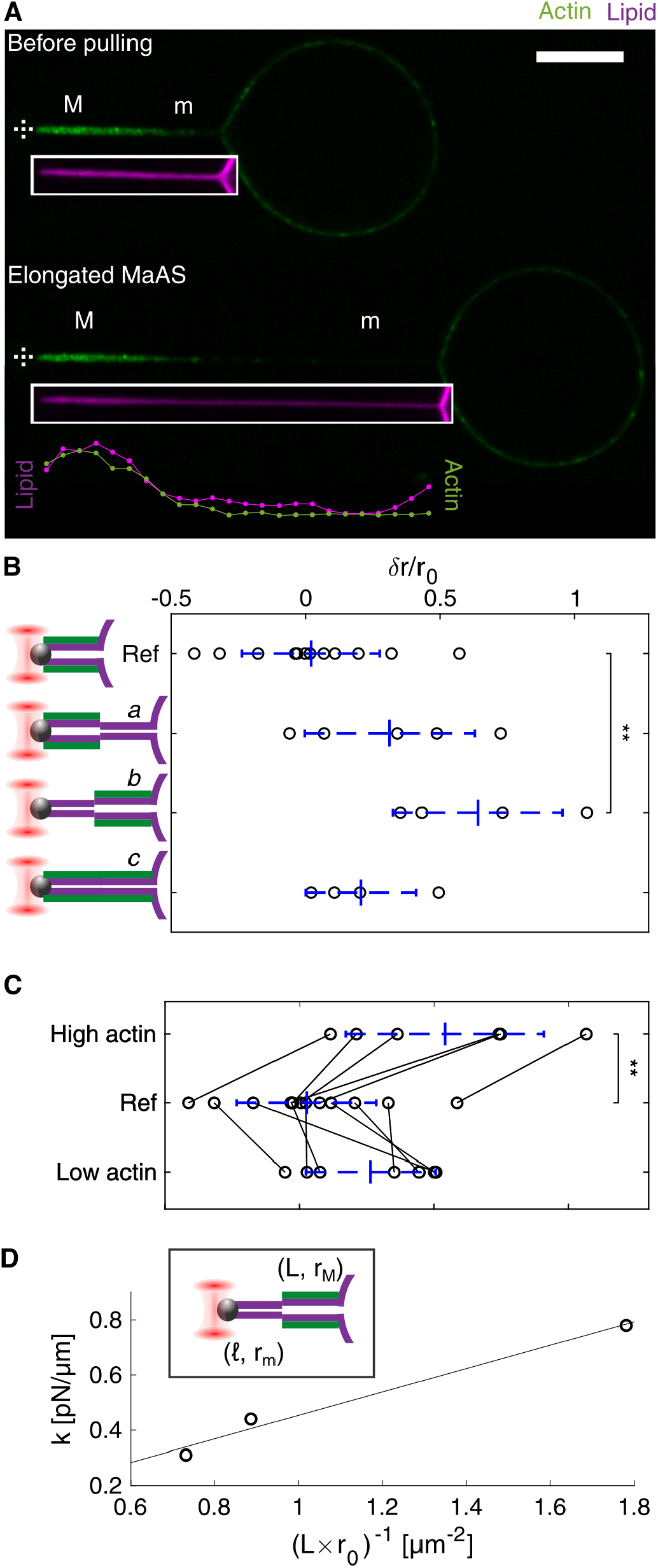
Membrane tube radius in elongated MaAS. (A) Representative confocal images before pulling for reference (top) and after elongation (bottom). Magenta and green correspond respectively to lipid and actin. The lipid image is displaced in the white rectangle right below the actin image for clarity. “M” and “m” regions are respectively defined as maximal and minimal actin intensity regions of the elongated MaAS (bottom) or before pulling (top). Graphs represent lipid and actin intensities along the elongated MaAS. Scale bar: 10 µm. Dashed crosses indicate bead center. (B, C) Relative difference in membrane tube radius between “M” and “m” regions defined in (A). (B) Values corresponding to classification schemed on the left (*a*, *b*, *c*). (C) “High actin” and “Low actin” refer respectively to actin content of elongated MaAS above and below average in Fig. 2A; reference condition (Ref) is before pulling. Lines connect same MaAS before and after elongation. (D) Proportionality factor of force-elongation curve, for *b*-types MaAS, as a function of (*L×r*_0_)^*−*1^. In the inset, parameters used for theoretical description of *b*-types MaAS. Data shown as mean + s.d. in (B, C). p-values calculated using the t-test. ** p *<* 0.01.

### Theoretical consideration of MaAS elongation

The increase in force necessary to obtain an elongated MaAS is proportional to its elongation (Fig. 2B) and the proportionality factor is slightly higher for *b*-types (Fig. S4A). This may be due to a hindrance of lipids flow under the actin sleeve that is higher in *b*-types than in *a* and *c* types. To account for this observation, we propose a model of force build-up assuming that lipids reaching the bare part of the tube are forced to go through the actin sleeve. As sketched in the inset in Fig. 3D, we model the bare section of membrane tube as a cylinder of a length *ℓ* and radius *r*_m_, that both vary over the course of tube extension, and the actin sleeve section has a fixed length *L* and a radius *r*_M_, that may also vary. The sleeve as well as the liposome are sheathed by actin, which binds to the membrane with an energy per area *W >* 0. This yields the following modified Helfrich energy function for the system (*17*)

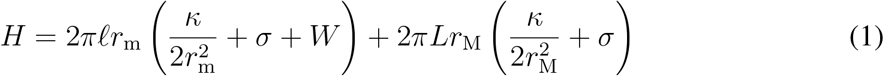

where *κ* is the bending rigidity of the membrane and *σ* the tension of the liposome.

Initially, the bead is in close contact with the actin sleeve, implying that *ℓ* = 0, from which we deduce 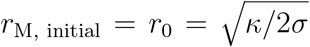 from the minimization of *H*. Assuming that the tube is pulled fast enough to prevent any substantial flow of lipids from the liposome to the MaAS, *ℓ* increases under the constraint of constant area *A* = 2*πr*_m_*ℓ* + 2*πr*_M_*L*. We thus minimize *H* with respect to *r*_m_ and *r*_M_ while fixing *A* for a given *ℓ* (supplementary materials). The resulting tube force reads *F* = d*H/*d*ℓ*. For small values of *W/σ* we thus obtain

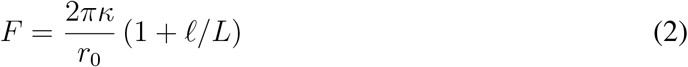

Therefore, *F* increases linearly with *ℓ*, in agreement with the observations of Fig. 2B upon tube pulling (supplemental materials). Equation (2) implies an apparent spring constant *k* = d*F/*d*ℓ* = 2*πκ/*(*r*_0_*L*), and fitting this relation with our experimental measurements leads to *κ* = 17*k*_B_*T* (Fig. 3D) close to the bare membrane value of 12*k*_B_*T*, where *k*_B_*T* is the thermal energy. We thus validate our assumption of lipids flow hindrance under the actin sleeve.

For the *a* and *c*-types configurations of Fig. 3B and S4A, the slope of the initial force-elongation is lower and close to the one of naked tubes, consistent with our assumption that the contact between the sleeve and the tube is largely responsible for limiting the lipid flow to the MaAS.

This theoretical description allows us to estimate the difference in radii between the actin sleeve and bare tube sections, through a calculation to next order in *W/σ*, yielding

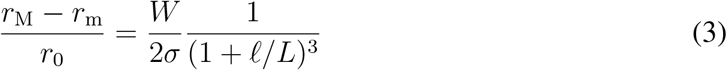

In practice this expression gives a good qualitative description of the *ℓ* dependence of the radius difference even for fairly large values of *W/σ ≃* 1 (supplementary materials). While comparing these predictions with our experimental measurements would in principle allow a determination of the actin binding energy to the membrane, we find in practice that these measurements do not yield a consistent value for *W/σ*, probably in part because the tension *σ* of the vesicle is not controlled in our experiment (supplementary materials). As a result, rather than determining the specific value of *W/σ*, in the following we place ourselves in the small-*W/σ* regime, which yields results qualitatively similar to the ones obtained at larger *W/σ* (Fig. S5).

### Relaxation of elongated MaAS

When maintained for several minutes, we observe that lipids fluorescence homogenizes along the tube as a function of time in elongated MaAS (dotted lines, Fig. 4A, B), more visible in cases *b* (Fig. 4B) where the heterogeneities are higher. Simultaneously, the force relaxes and mirrors radius relaxation (Fig. 4C, D). The characteristic time of force relaxation, estimated through an exponential fit, is longer for elongated MaAS than for naked tubes (Fig. 4E).

**Fig. 4.**
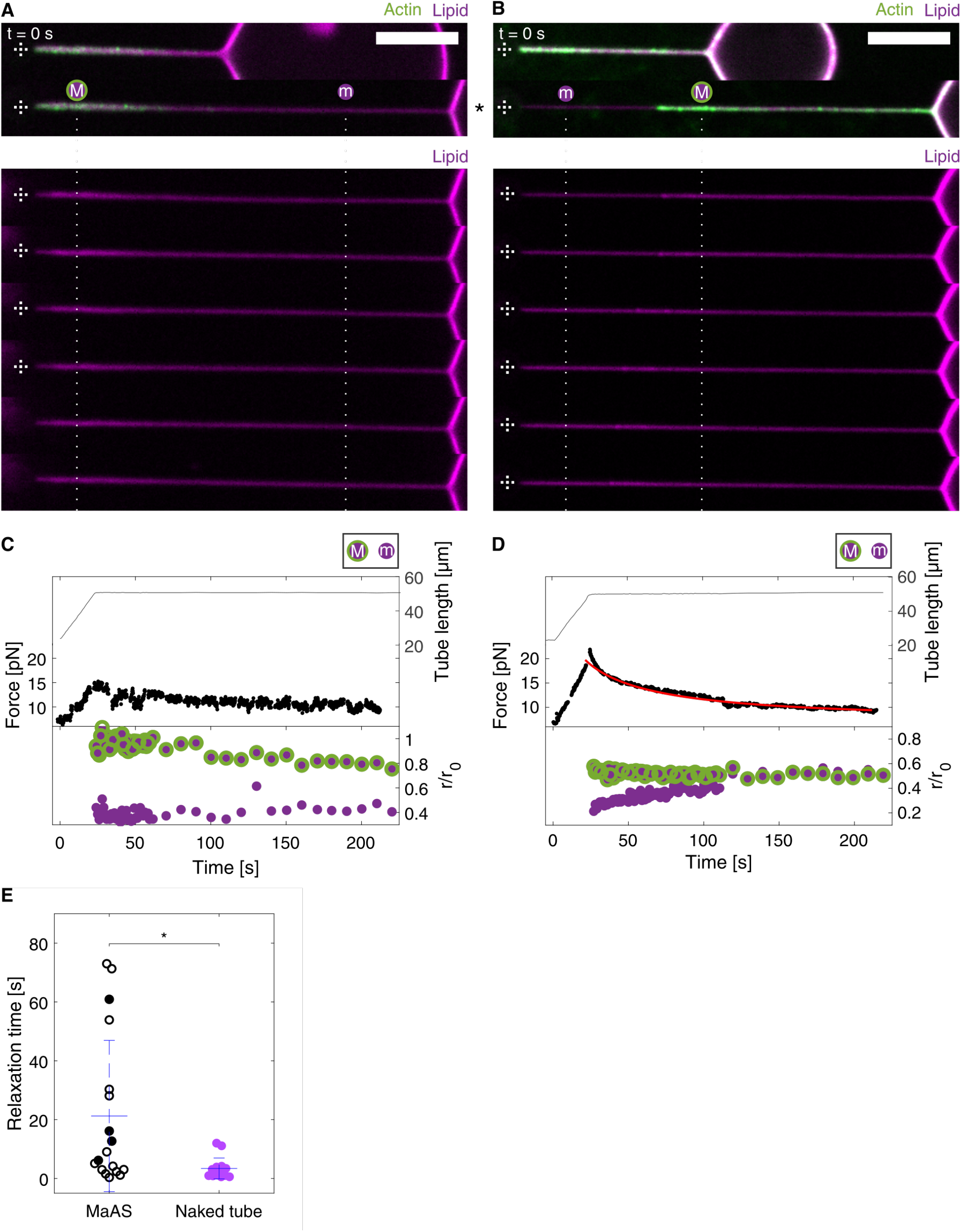
Elongated MaAS relaxation. (A, B) Representative confocal images of relaxing elongated MaAS (images every 30 s). Scale bars: 10 µm. Dashed crosses indicate bead center. Dotted lines follow lipid fluorescence relaxation. (C, D) Corresponding curves as a function of time for length, force and relative membrane tube radii associated with “m” and “M” regions defined in Fig. 3A and supplementary materials. Time 0 corresponds to the start of MaAS pulling. In red, the fitting curve from equation 4 using experimental values *ℓ* = 26.5 µm and *L* = 22.1 µm. (E) Relaxation time of the force for elongated MaAS and naked tubes, using an exponential decay (supplementary materials). Black filled circles point **b**-types MaAS. p-values calculated using the t-test. * p *<* 0.05.

During the initial rapid pulling step, the membrane becomes tenser and thinner than would be the case for a pure membrane tube. This results in the force increase of equation (2) for *b*-type. As the bead is held in position, relaxation occurs through lipids slowly flowing from the liposome towards the MaAS. By balancing the membrane forces driving this flow with the dissipation due to the friction of a membrane of viscosity *η* against a density *ρ* of actin attachment points, we compute the force relaxation dynamics to lowest order in *W/σ* as:

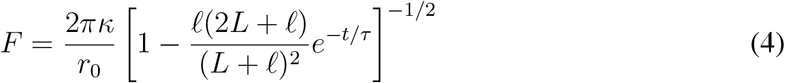

where the relaxation time is given by (supplementary materials):

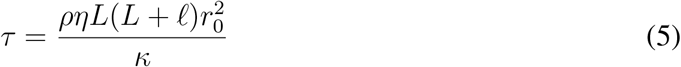

The fit of our data using equations (4) and (5) leads us to derive *ρη* = (3 *±* 1) *×* 10^6^ Pa ⋅ s/m (n = 4 *b*-types). By assuming we saturate all the binders, we can estimate in our experiment *ρ* = 10^15^ m^*−*2^ (*18*) which leads to *η* = 3 *±* 1 *×* 10^*−*9^ Pa ⋅ s ⋅ m close to previously published estimate *η* = 10^*−*8^ Pa ⋅ s ⋅ m (*19, 20*).

Therefore, elongated MaAS relaxation can be explained by the friction of lipids under the actin sleeve for *b* cases. For cases *c*, the friction is lower and therefore the relaxation time is much smaller, and close to the one of naked tubes.

### Elongated MaAS retraction

When the trap is turned off, we observe either partial or complete retraction (Fig. 5). Partial retraction ends up with the presence of an actin sleeve in between the bead and the liposome and its characteristic retraction time is 1.3 *±* 0.3 s (n = 10, Fig. 5A and open circles in Fig. 5C, D). Total retraction ends up with the bead at the liposome surface within a time that is smaller than 0.39 *±* 0.10 s (n = 7, crosses in Fig. 5C, D), similar to when pVCA is omitted (Fig. 5B and grey filled circles in Fig. 5C). Interestingly, regions of membrane tubes devoid of actin in elongated MaAS noticeably thicken when the MaAS retracts whereas they remain intact under the sleeve (arrow heads, Fig. 5A). Note that such a situation of a MaAS at zero force provides an even larger range of tube radii than when the tube is under force.

**Fig. 5.**
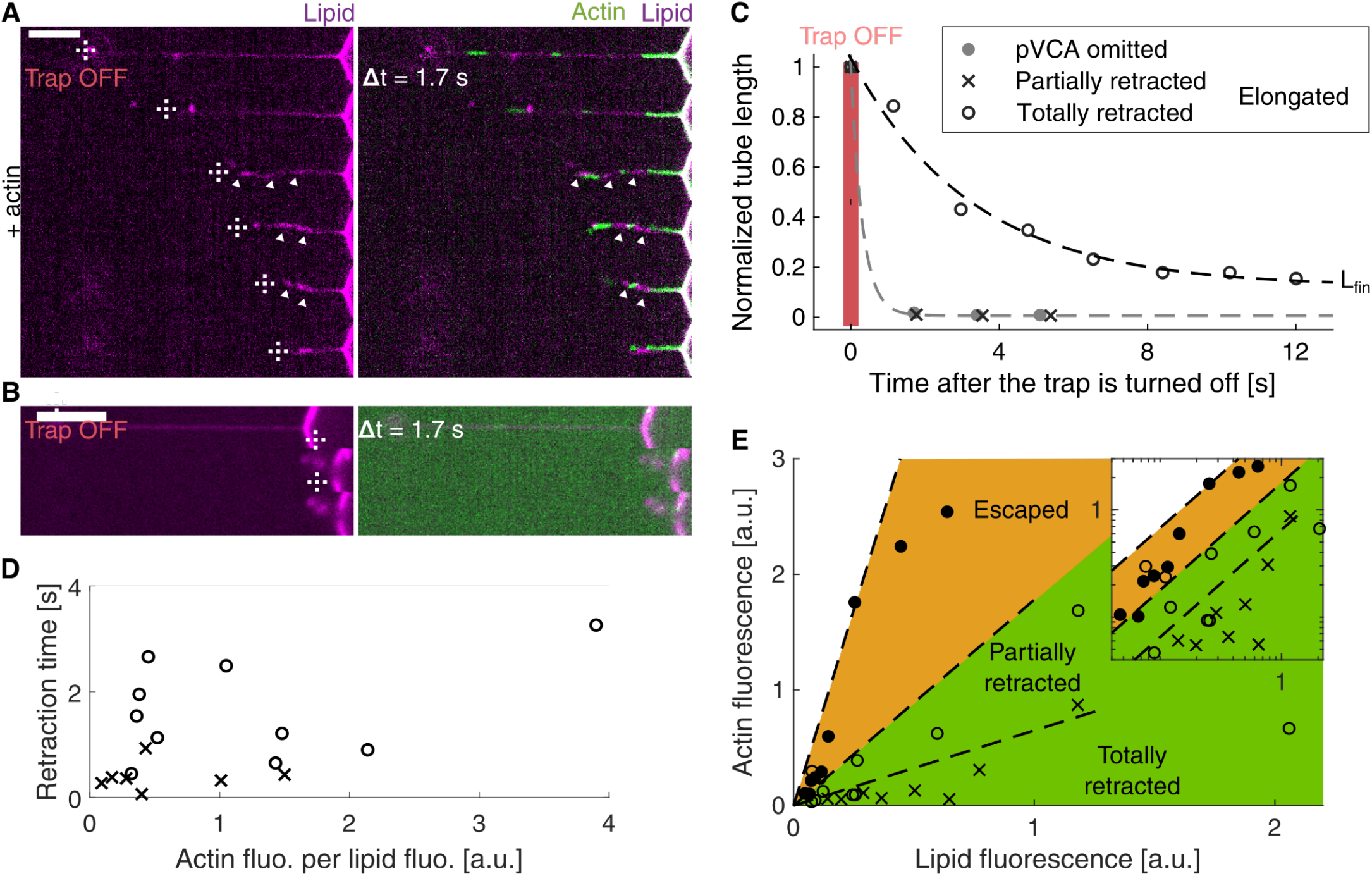
Elongated MaAS retraction. (A, B) Time lapse overlay images after the trap is turned off for an elongated MaAS (A) and control (pVCA microinjection is omitted) (B). Arrow heads indicate regions devoid of actin that get thicker during retraction. Scale: 10 µm. Dashed crosses indicate bead center. (C-E) Empty circle: partially retracted; crosses: totally retracted. (C) Tube length as a function of time after the trap is turned off for elongated MaAS or when pVCA is omitted (grey filled circles). (D) Retraction time as a function of actin content. (E) Actin fluorescence as a function of lipid fluorescence. Full circles are escaped MaAS. Dotted lines separate sectors. We use the orthogonal residue method, where we minimize the orthogonal distance between data and the linear regression to approximate sector separations. Elongated MaAS are in the green region, escaped MaAS are in the orange region. Totally and partially retracted MaAS are separated in sectors inside the green region. Inset: log-log representation, previous triangular sections then become bands here.

### Actin network in all MaAS types

All three MaAS types presented above are gathered in a diagram with their corresponding actin and lipid fluorescence intensities and reveal distinct regions depending on their types (see sectors or bands respectively in Fig. 5E or inset). We find that for a given tube radius, the amount of actin determines MaAS fate: escaped MaAS occur at the highest actin content that correspond to highly entangled networks (filled circles in Fig. 5E and Fig. 2C-top), and elongated MaAS separate in two sectors depending on their retraction fate (emptied circles and crosses in Fig. 5E). Note that thin tubes (lipid intensity low) cannot provide any sufficient support for the growth of an actin sleeve (empty sector zone top left corner, Fig. 5E), as their radius are too small to authorize correct building and growth of the actin network. Considering that we saturate nickel lipid sites with pVCA, the concentration of activators of polymerization at the surface of tubes is equal to the concentration of nickel lipids, and therefore proportional to the lipid intensity *I*_lipid_. Assuming that the quantity of actin *I*_actin_ in the sleeve is proportional to the quantity of activators the ratio *I*_actin_*/I*_lipid_ depends on the sleeve thickness and varies between sectors in Fig. 5E (or bands in inset) depending on the nature of MaAS (escaped, elongated, partial or total retraction).

Taking our estimate above for the sleeve radius in escaped MaAS (10 meshsizes, Fig. 2E) and matching it to the escaped MaAS sector (Fig. 5E), sleeve radius estimates for elongated MaAS can be inferred by their sector slopes. We find that elongated MaAS partially retracted display a sleeve of thickness in the 45 *−* 120 nm range (2-4 meshsizes), and that totally retracted elongated MaAS display actin sleeves of less than 45 nm (*∼* 2 meshsizes).

This experimental diagram summarizes that a sleeve thickness of 10 meshsizes is enough to stabilize membrane tubes, whereas 2 to 10 meshsizes only partially stabilize them. MaAS behave like naked tubes below 2 meshsizes. Therefore, stabilization of membrane tubes is highly sensitive to the number of meshes. Experiment and simulation highlight the importance of the cohesion of the the actin network around a membrane tube.

## Discussion

In summary, MaAS diversity can be explained by the thickness of their actin sleeve. Either they are totally cohesive by filament entanglement and their behaviour is controlled by the actin network elasticity, or they are extensible because actin filaments disentangle massively.

The role of actin in the morphology of intracellular membranes during trafficking or shaping the endoplasmic reticulum has been questioned in the last decade (*1, 4*). The impressive ability of actin dynamics to change rapidly the shape of membranes (*21–24*) naturally points to a similar role in membrane reorganization during trafficking. However, we show here unambiguously that a branched actin network grown through the Arp2/3 complex activated at the membrane surface has a stabilizing effect on the shape of the membrane tube rather than works as a scissionner.

Interestingly, the presence of a discontinuous actin sleeve allows some portions of tubes to get thinner under force whereas they get thicker when the force vanishes. Such a mechanism allows a membrane tube to provide portions of different radii along its length. This could explain the localization of specialized proteins such as BAR domains that detect curvature for scission (*25*) or dynamin that actively squeezes tubes in a curvature-dependent fashion (*26*). Moreover, portions of membrane tubes of different radii are connected by neck structures that may further serve for mechanisms of membrane reorganization.

Whereas the actin network meshsize is very close to the radius of tubular intermediates in trafficking and membrane tubes in our experiments, the number of meshes, above two, is what counts to obtain a membrane tube with a variety of radii. Note that a totally stabilizing network of 10 meshes is formed in few seconds, consistent with the characteristic time of tubular intermediates morphology changes in cells (*7*). An additional role of actin, demonstrated here for membrane tubes, is that it hinders lipid mobility, a mechanism proposed earlier to promote tube scission (*27*).

## Acknowledgments

This work was supported by the French Agence Nationale pour la Recherche (ANR), grants ANR 09BLAN0283, ANR 12BSV5001401, ANR 15CE13000403, and ANR 18CE13000701 by Fondation pour la Recherche Médicale, grant DEQ20120323737, and ERC Starting Grant 677532. Our groups belong to the CNRS consortium CellTiss. A.A performed experiments and analysed data. M.B. performed the simulations. M.L. and F.B.-W. developed the theoretical models. T.B., C.S., M.A.-G., J.L., F.V., K.G., J.P. and C.C. contributed to experimental data; J.M. purified the proteins; T.B., C.C. and C.S. designed the research. All authors contributed to writing the paper. The authors declare no competing interests. The data that support the plots within this paper and other findings of this study are available from the corresponding author upon request.

## Supplementary materials

### Materials and Methods

#### Buffer solutions

Chemicals are purchased from Sigma Aldrich unless specified otherwise. The internal buffer (TPI) consists in 2 mM Tris and 200 mM sucrose. The external buffer (TPE), where the polymerization occurs, contains 1 mM Tris, 50 mM KCl, 2 mM MgCl_2_, 0.1 mM DTT, 2 mM ATP, 0.02 g/L *β*-casein and 95 mM sucrose. TPE_inj_, limiting actin polymerization inside the micropipette, consists in 1 mM Tris, 1 mM MgCl_2_, 0.1 mM DTT, 0.02 g/L *β*-casein and 195 mM sucrose. TPA, a high osmolarity buffer, contains 1 mM Tris, 1 mM MgCl_2_, 0.1 mM DTT, 0.02 g/L *β*-casein and 395 mM sucrose. All buffers are adjusted at pH 7.4 and their osmolarity is set at 200 mosm/kg (400 mosm/kg for TPA). Osmolarities are measured with a vapor pressure osmometer (Vapro 5600, Wescor, USA). G-buffer, to obtain monomeric actin, is composed of 2 mM Tris, 0.2 mM CaCl_2_, 0.2 mM DTT, 0.2 mM ATP (pH 8.0).

#### Preparation of liposomes

Lipids stocks EPC (L-*α*-phosphatidylcholine from egg yolk), DS- PE-PEG(2000)-biotin (1,2-distearoyl-sn-glycero-3-phosphoethanolamine-N [biotinyl-(polyethylene glycol) 200]) and 18:1 DGS-NTA(Ni) (1,2-dioleoyl-sn-glycero-3-[(N-(5-amino-1-carboxypentyl) iminodiacetic acid)succinyl]) are purchased from Avanti Polar Lipids (Alabaster, USA). Texas Red DHPE (1,2-dihexadecanoyl-sn-glycero-3-phosphoethanolamine, triethylammonium salt) is obtained from Thermo Fisher (Waltham, USA). All stocks are aliquoted in chloroform/methanol at volume ratio 5/3.

Liposomes are formed using the standard electroformation method (*28*). The lipid mix (molar ratio EPC/DGS-Ni/DSPE-PEG-biotin/Texas Red DHPE of 89.4/10/0.1/0.5) is dissolved at 2.5 g/L in chloroform/methanol at volume ratio 5/3. A volume of 5 µL of this solution is spread on an ITO-coated (Indium Tin Oxide) glass slide (63691610PAK, Sigma Aldrich, Germany). The film of lipids is dried in vacuum for 2 h. Then the films of lipid are hydrated with TPI by assembling the two conductive slides facing each other into a chamber sealed with Vitrex (Vitrex Medical A/S, Denmark). An oscillating electric field (10 Hz, 3 V peak to peak) is applied across the chamber during 2 h. Liposomes are stored at 4°C for up to two weeks.

#### Proteins and reagents

Actin and porcine Arp2/3 complex are purchased from Cytoskeleton (Denver, USA). Fluorescent Alexa Fluor 488 actin conjugate (actin-488) is purchased from Molecular Probes (Eugene, USA). Mouse *α*1*β*2 capping protein (CP) is purified as described elsewhere (*29*). His-pVCA-GST (pVCA, the proline rich domain-verprolin homology-central-acidic sequence from human WASP, starting at amino acid Gln150) is purified as for PRD-VCAWAVE (*30*). Untagged human profilin is purified as in (*22*). A solution of 30 µM monomeric actin containing 15% of labelled actin-488 is obtained by incubating the actin solution in G-Buffer over two days at 4°C. Commercial proteins are used with no further purification and all concentrations are checked by a Bradford assay.

#### Optical tweezers, image acquisition and tube pulling

As previously described (*31*) we use a system that allows us to simultaneously measure forces with optical tweezers and record images with a spinning disk confocal microscope (CSUX1 YOKOGAWA, Andor Technology, Ireland) and by a high resolution sCMOS Camera (Andor Neo, Ireland). The optical tweezers are based on an infrared laser (*λ* = 1064 nm, *P* = 5 W, YLM-5-LP-SC, IPG Laser, Germany) controlled by an XY AOD pair (MT80-A1 51064 nm, AA Opto Electronic, France). A water-immersion objective (PLAN APO VC 60x A/1.2WI IFN 25 DIC N2, Nikon, Japan) is used for imagery.

To pull a membrane tube, a streptavidin-coated polystyrene bead (3.05 µm diameter, strepta-vidin-coated, Spherotech, Illinois, USA), which serves as a handle to maintain the tube, is first trapped optically. A biotinylated liposome, slightly adherent to the bottom surface of the chamber, is then attached to the bead. By moving away the stage at a constant speed, a tube forms between the liposome and the bead. The position of the bead relative to trap center is recorded based on the back focal plane technique (*32*). We record the interference signal between the unscattered laser light and the light scattered by the bead, imaged on a Quadrant PhotoDiode (QPD, PDQ-30-C, Thorlabs, Germany) placed on the optical path. After proper calibration, the voltage from the QPD is proportional to the bead displacement. Trap stiffness *k*_trap_ is deduced from the power spectrum of the bead fluctuations around the center of the trap. We find *k*_trap_ = 58.4 ± 2.3 pN/µm over 25 independent experiments. Altogether, calibrations of the QPD and the trap stiffness provide force measurements from the bead displacement in the trap.

Finally, the chamber is mounted on a 2D piezo stage (MS 2000, ASI, USA) that controls its positioning and allows its displacement at a controlled velocity. The tube elongation *∆ℓ* is experimentally calculated from the known speed *v* of the stage and the position *x* of the bead with respect to the trap center. Then, during elongation *vt* = *∆ℓ* + *∆x* where *∆x* = *∆F*_QPD_*/k*_trap_ is the relative bead displacement in the trap. Finally, tube elongation yields *∆ℓ* = *vt − ∆F*_QPD_*/k*_trap_.

#### Chamber and micropipette preparation

Prior to experiments, we sonicate glass coverslips (0.13 − 0.16 mm, Menzel Gläze, Australia) in 2-propanol for 5 minutes, extensively rinsed with water and dried under filtrated compressed air. Then the glass surface is activated by a plasma cleaner (PDC-32G, Harrick Plasma, USA) during 2 minutes, followed by a 30 minutes passivation using 0.1 g/L PLL(20)-g[3.5]-PEG(2) (SuSos, Switzerland) in a 10 mM Hepes solution (pH 7.4). The experimental chamber is made of two glass coverslips separated by a 1 mm steel spacer. The chamber is filled with a 100 µL polymerization mix (P-solution) in TPE containing 3 µM profilin, 37 nM Arp2/3 complex, 25 nM CP, 2 µL liposomes in TPI, and 1 µL polystyrene beads diluted 100 times in TPE.

Micropipettes are prepared from borosilicate capillaries (0.7mm/1.0 mm for inner/outer diameter, Harvard Apparatus, USA), using a puller (P2000, Sutter Instrument, USA) with parameters previously described in (*31*). Micropipette tips are then micro-forged (MF 830, Narishig, Japan) up to an internal diameter of 10 µm. Micropipettes are filled by aspirating 1 µL of the desired solution. Mineral oil is filled on the other side of the micropipette using a MicroFil (250 µm ID 350 µm OD 97 mm long, World Precision Instrument, UK). We prepare two micropipettes: the first one contains 2 µM pVCA, 0.01 g/L sulforhodamine-B (to monitor the microinjection), in TPE; the second one contains 3 µM actin-488 and 3 µM profilin, in TPE_inj_, adjusted to the osmolarity of 200 mOsm/kg with TPA.

Note that profilin is present in the actin microinjection pipette and in the P-solution, so that actin polymerization is prevented in the micropipette and in solution, and occurs only at the membrane surface. Each micropipette is set up into the chamber, one on each side connected to two separated reservoirs to control the injection pressure. The chamber is sealed on each side by adding mineral oil, to block evaporation over the time of the experiment.

#### Fluorescence intensity

We define a box centered around the tube, the height of which is kept at 50 pixels corresponding to 6.9 µm (Fig. S3A-C). The background fluorescence per pixel along the *x* axis is taken on the first pixel row at the top of the box and is subtracted to each box pixel. We then determine the intensity of lipid, *i*_lip_(*x*), and the intensity of actin, *i*_act_(*x*) as the total fluorescence intensity along the vertical *y* axis (Fig. S3B). The actin (resp. lipid) fluorescence is then defined as the average of the intensity along the variable *x*, 〈*i*_act_(*x*)〉_*x*_ (resp. 〈*i*_lip_(*x*)〉_*x*_). The actin fluorescence per lipid fluorescence is determined as the ratio 〈*i*_act_(*x*)〉_*x*_/〈*i*_lip_(*x*)〉_*x*_. Local intensities of actin or lipid (respectively *I*_act_(*X*) and *I*_lip_(*X*)) are sliding averages as defined in Fig. S3C.

The membrane tube radius is proportional to the number of fluorescently labeled lipids as described in (*33*) and shown in Fig. S3D. To quantify heterogeneities along a membrane tube, after elongation, we define relative variation of radii along a tube, compared to the initial radius, as the relative variation of lipid fluorescence, compared to the initial lipid fluorescence: 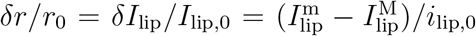, where the superscripts m and M respectively refer to region where actin intensity is minimal (resp. maximal). When the lipid tube is homogeneous, *δI*_lip_ is centered around zero but can also have negative values due to our definition of m and M from the actin signal (Fig. S3A-C).

#### Characteristic relaxation and retraction times

At the end of elongation, force relaxation is fitted by a first order decreasing exponential: *F* (*t*) = *A* exp (− *t/τ*_rel_) + *F*_fin_, where *F*_fin_ is the final value of the force. The maximal force after elongation is given by *F*_max_ = *A* + *F*_fin_.

During retraction after the laser is turned off, we determine the tube length *ℓ* over time *t* by manual tracking using ImageJ. The characteristic retraction time *τ*_ret_ is determined by the fit *L*(*t*) = *a* exp (− *t/τ*_ret_) + *L*_fin_. We define that there is partial retraction if *L*_fin_ is higher than 2 µm.

#### Statistical analysis

Results are represented as mean standard error mean (s.e.m.). Graphs show the mean standard deviation (s.d.). All statistical analyses are performed using MedCalc software. A *t*-test is used to determine the statistical significance and p-values are indicated (n.s., non significant; *p *<* 0.05, **p *<* 0.01, ***p *<* 0.001).

## Supplementary Text

### 1 Simulation

#### A microscopic model for branched actin networks

##### Growth algorithm

We first design the branched actin structure through a lattice-based growth model. In our model, filaments occupy the nodes of a three-dimensional regular (e.g., body centered cubic or BCC) lattice and grow unidirectionally, in one of the sixth *ϵ*_*x*_**x** + *ϵ*_**y**_**y** + **z** directions, with *ϵ*_*x,y*_ ∈ {1, 0, −1}. Filaments grow longitudinally and branching happens with a probability *β* that sets the concentration of Arp2/3. It dictates the relationship between meshsize and thickness of the filaments *d* through 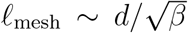. Its exact value does not matter in the limit of thin filaments where *β* is small. We choose *β* = 0.03 which in practice is a good balance between accuracy in smallness and the computational time. To prevent nonphysical, exponentially growing actin densities, we impose that there can be only one filament per lattice site. After a transient regime, this process leads to an intertwined branched actin network of dimension shown in Fig. S6A where the density of the structure reach a constant. A cubic portion of the grown system of dimension 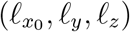 is cut (in the stationary regime), discretized into monomers and will be used as initial configuration for our simulations.

##### Coarse-grained numerical model

We developed a minimal coarse grained model for branched actin networks. The actin filaments of the grown structure shown in Fig. S6A are represented by a series of discrete monomer beads of diameter *d* (Fig. S6B) that interact through a potential composed of three terms:

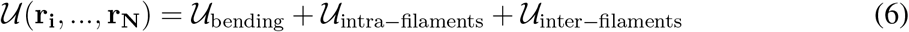

The first term 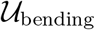 is a three body interaction that accounts for the bending rigidity of the filaments. It is given by:

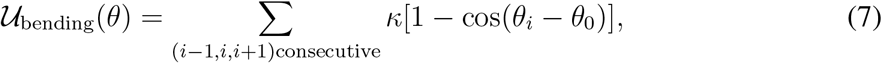

where *κ* sets the energy scale for the resistance to bending and is related to the persistence length of the filaments via *ℓ*_*p*_ = *κd/k*_B_*T*. Here we will explore the athermal limit that correspond to semi-flexible filaments. The angle *θ* is given by a triplet of three consecutive monomers (Fig. S6B). The equilibrium angle *θ*_0_ is set equal to *θ*_0_ = 180° for all triplets to have straight filaments under no stress, however different values are used to account for the specific microscopic morphological structure of the branched actin, which are characterized by three angles at the branching point equal to: 70°, 110°and 180°(Fig. S6B left panel).

The second contribution in 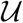 bind monomers belonging to the same filament together through a finitely extensible nonlinear elastic (FENE) potential coupled to a repulsive Lennard-Jones term preventing monomers to overlap:

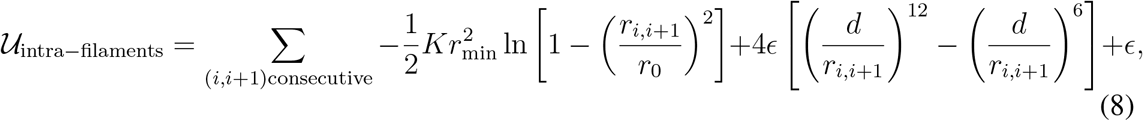

with **r**_i,i+1_ = **r**_i+1_ − r_i_, with **r**_i_ denoting the position vector of the *i*-th monomer in the filament, *r*_0_ is the maximum bond length extension, *K* the bonding energy scale and *ϵ* the strength of the repulsive part.

Finally the last contribution is a truncated and shifted Lennard-Jones potential of form:

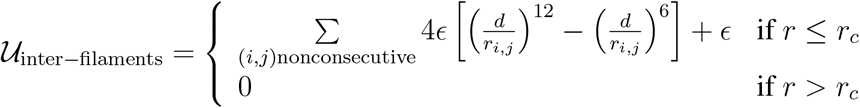

This term acts as an excluded volume between filaments with *r*_*c*_ the cut-off distance chosen equal to the minimum of the Lennard-Jones potential *r*_*c*_ = 2^1/6^*d* (Fig. S6B right panel). With respect to the mass unit *m* = 1, length unit set by the monomer diameter *d* = 1, and the energy scale given by *κ* = 1, we set *r*_min_ = 2, *ϵ* = 1, *K* = 15.

##### Relaxation of the structure

With this model, we have performed molecular dynamics simulations. The simplicity of the model allows us to run large scale simulations with up to *N* ≃ 10^5^ monomers, which is essential for a statistical analysis of the microscopic dynamics. The initial configuration composed of *N* monomer in a simulation box of size 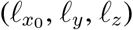 is prepared with the protocol which consists in (i) starting from a grown structure configuration, (ii) turning on the interactions and (iii) letting the network reach mechanical equilibrium where the residual stresses are relaxed by monitoring the box size through a barostat. Once the pressure vanishes, the kinetic energy is then completely drawn from the system by means of a dissipative microscopic dynamics:

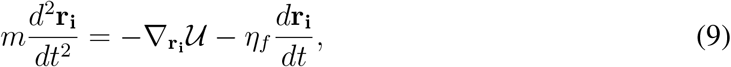

Here *η*_*f*_ is the damping coefficient associated with coupling of the monomer motion to the surrounding fluid. All simulations are performed in the over-damped limit which correspond to *η*_*f*_ = 1. The time step *δt*_0_ used for the numerical integration is *δt*_0_ = 0.005.

All simulations discussed here have been performed using a version of LAMMPS (*34*).

##### Uniaxial deformation

To determine the network mechanical response, each monomer configuration can be submitted to a series of incremental strain steps (*35, 36*). In each step we increase the cumulative deformation strain by an amount 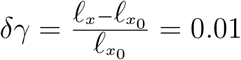 by first applying an instantaneous linear deformation *Γ*_*δγ*_, corresponding to simple uniaxial deformation in the *x* axis, to all monomers:

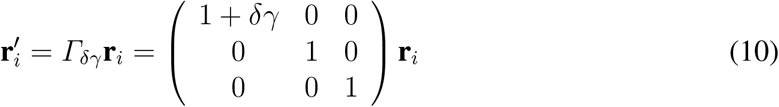

The Lees-Edwards boundary conditions are updated as well, to comply with the increase in the cumulative strain. The configuration 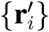 is no longer a minimum of the potential energy, and the small deformation step induces unbalanced internal forces. Hence we relax the linearly deformed configuration by letting the system free to evolve in time while keeping the global strain constant:

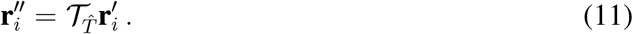

where 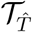 is the time evolution operator for a specified time interval 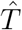 and given by the damped dynamics 9.

After *n* steps, the cumulative strain is *γ*_*n*_ = *n δγ* and the network configuration is

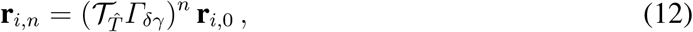

where {**r**_*i*,0_} denotes the configuration of the starting inherent structure.

In this procedure instead of using an energy minimization algorithm after each linear deformation step, we follow the natural dynamics of the system (with a viscous energy dissipation) for a prescribed time interval 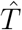. We can therefore define a finite rate of deformation 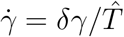 We probe the quasi-static limit for which 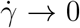 and is sufficiently small so that our results do not depend on its specific value (Fig. S6C). Disregarding effects due to monomer inertia, the microscopic dynamics 9 introduce a natural time scale *τ*_0_ = *η_f_ d*^2^*/κ*, corresponding to the time it takes a monomer subjected to a typical force *κ/σ* to move a distance equal to its size. To mimic the cohesion of the actin filaments on the surface of the membrane we fix the *z* position of the monomer at the top of the box where *z* = *ℓ*_*z*_ and let them free to move in the *xy* plane.

##### Stress and force calculation

The average state of stress of the network is given by the virial stresses as 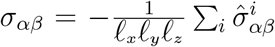, where the Greek subscripts stand for the Cartesian components *x, y, z*, (*ℓ_x_, ℓ_y_, ℓ*_*z*_) the size of the system and 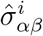 represents the contribution to the stress tensor of all the interactions involving the monomer *i* (*37, 38*). The latter contribution is calculated for each monomer, by splitting the contributions of the two-body and the three-body forces according to the following equation:

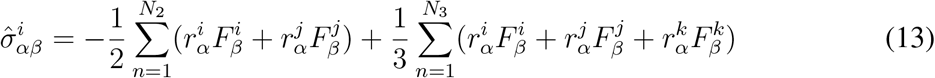

The first term denotes the contribution of the two-body interaction, where the sum runs over all the *N*_2_ pairs of interactions that involve the monomer *i*. The couples (*r*^*i*^, *F*^*i*^) and (*r*^*j*^, *F*^*j*^) denote respectively the position and the forces on the two interacting monomers. In the same way, the second term indicates the three-body interactions (bending) involving the monomer *i* and two neighbors denoted by the label *j* and *k*.

Note that for overdamped athermal semi-flexible actin networks in the quasi-static limit, the kinetic contribution 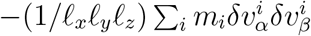 to the global stress tensor (from the fluctuations of the monomer velocities with respect to the average) is negligible. Under uniaxial deformation, the average force *F* is then given by the average stress *σ*_*xx*_ extracted from the simulations times the cross-section of the network. To derive the corresponding force for the whole cylinder actin sleeve of inner radius *r*_0_ = 25 nm and variable thickness *ℓ*_*z*_, the cross-section is given by 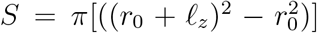. The force unit is our simulation has a unit of *κ/d*, since the meshsize is given by 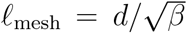, this lead to 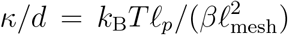. To convert in SI units we haven chosen *k_B_T* = 4.10^*−*21^*J*, *ℓ*_*p*_ = 10 µm (*15*) and *ℓ*_mesh_ = 30 nm (*16*). All our results are taken in the quasi-static limit. Fig. S6C shows force-deformation curves for different rates of deformation. This limit in reached for sufficiently slow relaxation that correspond to 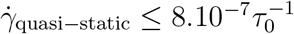.

## 2 Theory

Here we present the detailed derivation of the results presented in the main text for the elongated MaAS pulling force, effective spring constant and the radii of the coated and naked sections of the MaAS.

We start from the modified Helfrich free energy detailed in the main text, which assumes that a membrane reservoir with tension *σ* covered by actin is in contact with a membrane tube of bending rigidity *κ* and length *L*+*ℓ* which is coated by actin over a length *L*. Over this length, the interactions between the actin and the membrane is modeled by a binding energy per unit surface *W*, yielding

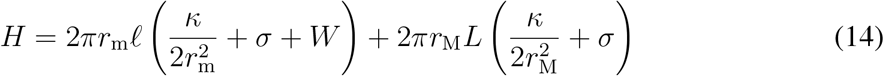

where *r*_M_ and *r*_m_ denote the radii of the coated and naked sections of the tube, respectively. In equation (14), the terms proportional to *κ* account for the bending energies of two cylinders of radii *r*_m_ and *r*_M_, respectively. To rationalize the form of the other terms, the transfer of a unit of surface of membrane from the actin-covered reservoir to the actin-free naked tube comes at a cost that is the sum of the work performed against the surface tension of the membrane and the unbinding of that unit of surface from the actin, hence the factor *σ* + *W* in the first term of the right-hand-side of equation (14). Conversely, the transfer of membrane from the reservoir to the coated section of the tube only involves the cost associated with working against the membrane tension. Indeed, while some membrane-actin bonds are broken when the membrane leaves the reservoir, an equal number is reformed as it joins the coated tube, implying that there is no net change in binding energy. Here we work in the ensemble where the lengths *L* and *ℓ* are fixed, implying that the often-introduced free energy contribution *F* (*L*+*ℓ*) accounting for the action of an external tube pulling force *F* would only contribute an irrelevant constant. Prior to the extension of the MaAS (*i.e.*, for *ℓ* = 0), minimizing this expression with respect to *r*_M_ yields a coated tube equilibrium radius 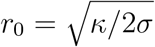.

Here we first study the changes to this initial state during the extension of a type-*b* elongated MaAS in Sec. 2. As the MaAS is being pulled relatively quickly, we assume that the lipid exchange with the reservoir over the duration of this step is negligible, although membrane flows *within* the MaAS reach a fast equilibrium. Section 2 then discusses the subsequent relaxation of the MaAS as the length of the tube is maintained over longer time scales, allowing the lipids to flow from the reservoir into the MaAS. This separation of time scales makes the implicit assumption that the internal relaxation of the MaAS is much faster than its equilibration with the membrane. While approximate, this approach yields a good agreement with experimental measurements as shown in the main text, suggesting that it captures the essential features of the MaAS relaxation.

### MaAS radii and force during fast pulling

The rapid extension of the type-*b* elongated MaAS implies the absence of lipid flow from the reservoir and the conservation of its area

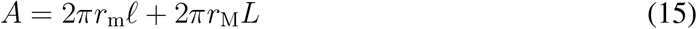

The dimensionless membrane area *a* = *A/*(2*πr*_0_*L*) thus remains equal to its initial value throughout the MaAS extension. For an initially equilibrated MaAS, this initial value is *a* = 1. In this section we however discuss the more general case of an arbitrary but constant value of *a*. This slight generalization will prove useful in Sec. 2, where lipid flows imply a change of the MaAS area and an increase of *a*. In this general case, the constraint of fixed area reads

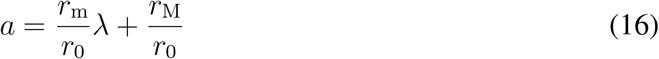

where we have defined *λ* = *ℓ/L*.

Although the total area of the MaAS is fixed, we consider that lipids can freely balance *within* the MaAS, implying that the system adopts the most energetically favorable values of *r*_m_ and *r*_M_ compatible with the constraint of equation (16). This constraint is automatically satisfied if we write the two radii as functions of a single free variable *x* as

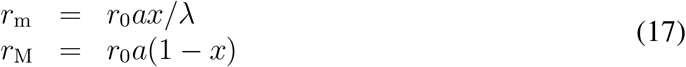

Introducing the dimensionless Hamiltonian 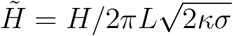 and the dimensionless binding energy *w* = *W/σ*, this implies that the equilibrium configuration of the MaAS is determined by minimizing

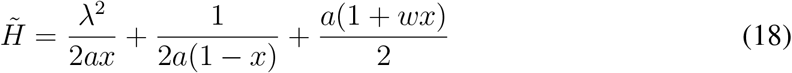

with respect to *x*. The two first terms in the right-hand side of equation (18) account for the bending energies of the naked and coated sections of the MaAS respectively. The third term described the binding energy within the coated region, and the surface tension terms are grouped within the fourth term. Minimizing with boils down to solving the following equation:

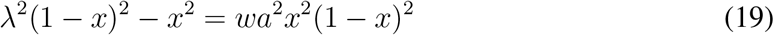

As all fourth-order polynomial equations, this equation has an explicit analytical solution, although a rather cumbersome one (Fig. S5).

Combining equation (19) with equations (17) for *a* = 1 allows to infer the value of the radii *r*_m_ and *r*_M_ at the end of the elongation phase. Taking our experimental values leads to *w* ≃ (17 ± 10) (n = 4 *b*-types). We find in practice that these measurements do not yield a consistent value for *w*, probably in part because the tension *σ* of the vesicle is not controlled in our experiment. As a result, in the following we place ourselves in the small-*w* regime, which yields results qualitatively similar to the ones obtained at larger *w*, with the added advantage of yielding expressions that are much more readable than the full solution of equation (18).

The most energetically favorable value of *x* thus reads

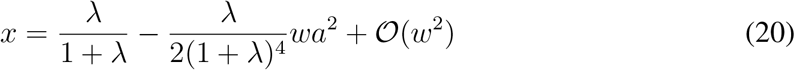

which we then use to derive the dimensionless versions of two quantities measured in the main text, namely the tube force and difference between the two MaAS radii:

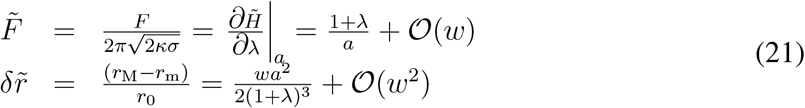

which yield equations (2) and (3) of the main text upon setting *a* = 1 as is appropriate for the initial extension of the MaAS. The linear dependence of 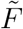 on *λ* in the small-*w* expression of equation (21) is in good agreement with the form of the initial force increase observed in Fig. 4C, D of the main text. Equation (21) additionally implies that

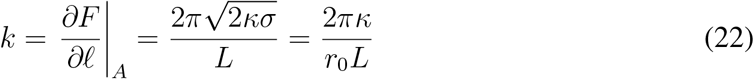

which we use to infer *r*_0_ from the fit of Fig. 3D.

### Membrane flow and force relaxation

In our experiments the length *L* + *ℓ* of the MaAS is held fixed following the elongation step discussed in Sec. 2. As this length is maintained over long time scales, lipids flow from the reservoir, driven by the difference in membrane chemical potential between the MaAS and the reservoir.

We denote this difference in membrane chemical potential per unit area as *µ* = d*H/*d*A*, where the total derivative indicates that *H* must be minimized with respect to *x* prior to the differentiation with respect to *A*. This minimization reflects our approximation that the internal degree of freedom *x* relaxes quickly compared to the time scales involved in lipid flow and therefore compared to any change in the value of *A*. Combining equations (18) and (20), we find

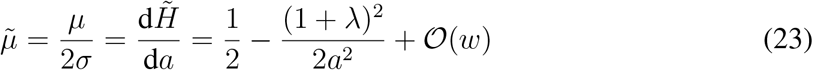

To compute the dynamics of the lipid flow, we assume that the associated dissipation stems primarily from the friction between the flowing lipids and the stationary actin in the coated section of the MaAS. Denoting by *ρ* the surface density of the actin attachment points onto the membrane, by *η* the surface viscosity of the membrane and by *v* the velocity of the lipids, the friction stress within the coated section of the MaAS is given by Darcy’s law as *Σ* = *ρηv*, implying a dissipated power *P*_*d*_ = 2*πr*_M_*Σv*. This dissipated power equals the power liberated by the flow of lipids from the reservoir to the MaAS, which by definition of the chemical potential difference *µ* reads *P*_*l*_ = *µ*d*A/*d*t*. Equating *P*_*d*_ and *P*_*l*_ while noting that d*A/*d*t* = 2*πr*_M_*v*, we find that the MaAS area is governed by the following evolution equation:

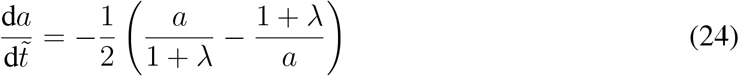

where we have defined the dimensionless time as

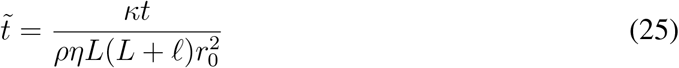

yielding the definition of *τ* in equation (5) of the main text. Imposing the initial condition 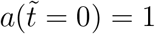 on equation (24) yields

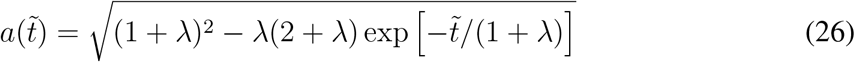

which we combine with equation (20) to derive the force relaxation law of equation (4) of the main text.

**Fig. S1.**
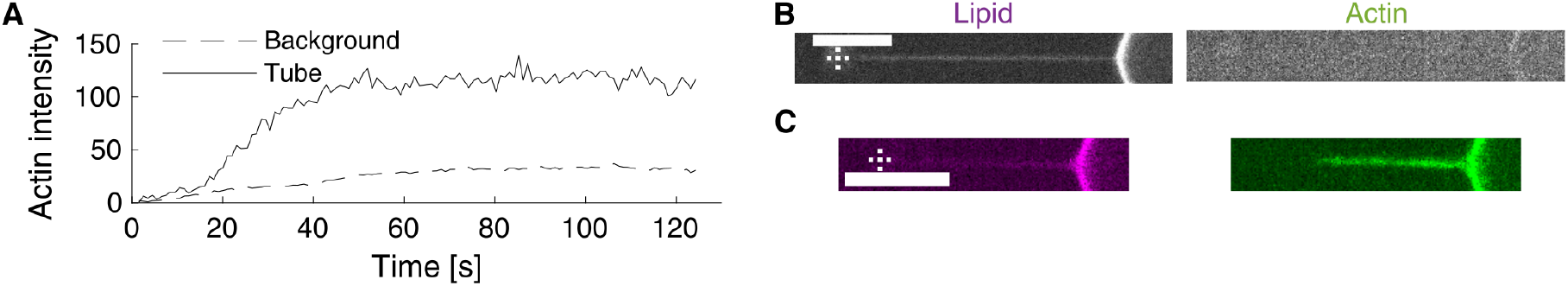
Recruitment of actin to a membrane nanotube. (A) Actin background, around the membrane nanotube, as a function of time shows that actin injection is on going. Actin intensity along the tube show specific polymerization of actin along the tube, that reaches a plateau after few tens of seconds. These data are extracted from the experiment described in Fig. 1A. (B) Absence of actin when pVCA is omitted. (C) Polymerization of an actin network on a preformed membrane tube activated by pVCA. Scale = 10 µm. Dashed crosses indicate bead center.

**Fig. S2.**
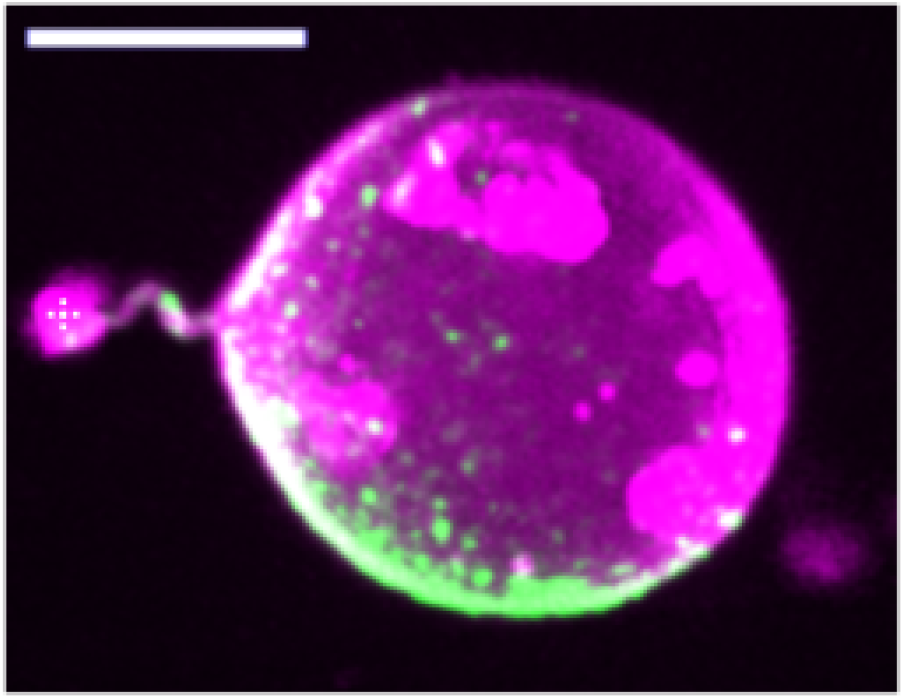
3D imaging of an escaped MaAS. 3D reconstruction of a MaAS imaged along Z-axis, 20 minutes after escape. The bead is connected to the MaAS at the left. Magenta and green correspond respectively to lipids and actin. Scale: 10 µm. Dashed cross indicates bead center.

**Fig. S3.**
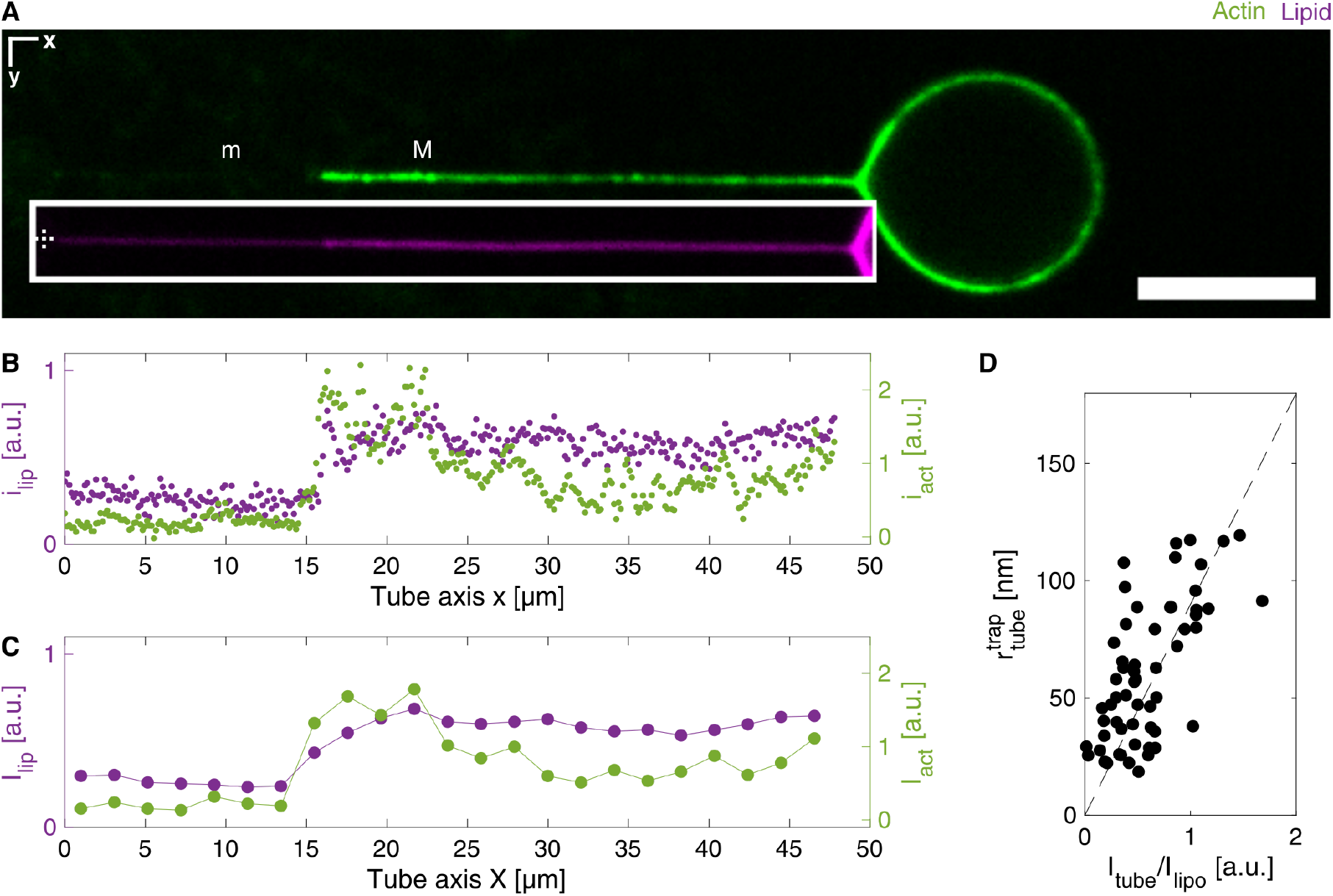
Fluorescence measurement method. (A) Representative spinning disk confocal images of MaAS. Magenta and green correspond respectively to lipids and actin. The lipid image is displaced in the white rectangle right below the actin image for clarity. *x* and *y* refer to horizontal and vertical axis. Scale: 10 µm. (B) *y*-integrated intensities *i*_lip_ and *i*_act_ as a function of *x*. (C) Sliding average of the intensities in (B), with a window of 2 µm, allow to define “m” and “M” regions in (A) (see supplementary materials). (D) Calibration of tube radius as a function the ratio between tube and vesicle intensities, as previously described in (*33*). We use the trap force *F* ^trap^ to calculate the corresponding radius 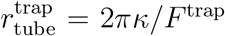, where *κ* = 12*k*_B_*T* is the bending modulus experimentally determined. This leads to *r*_tube_ = (90 *±* 10 nm)*i*_tube_*/i*_lipo_.

**Fig. S4.**
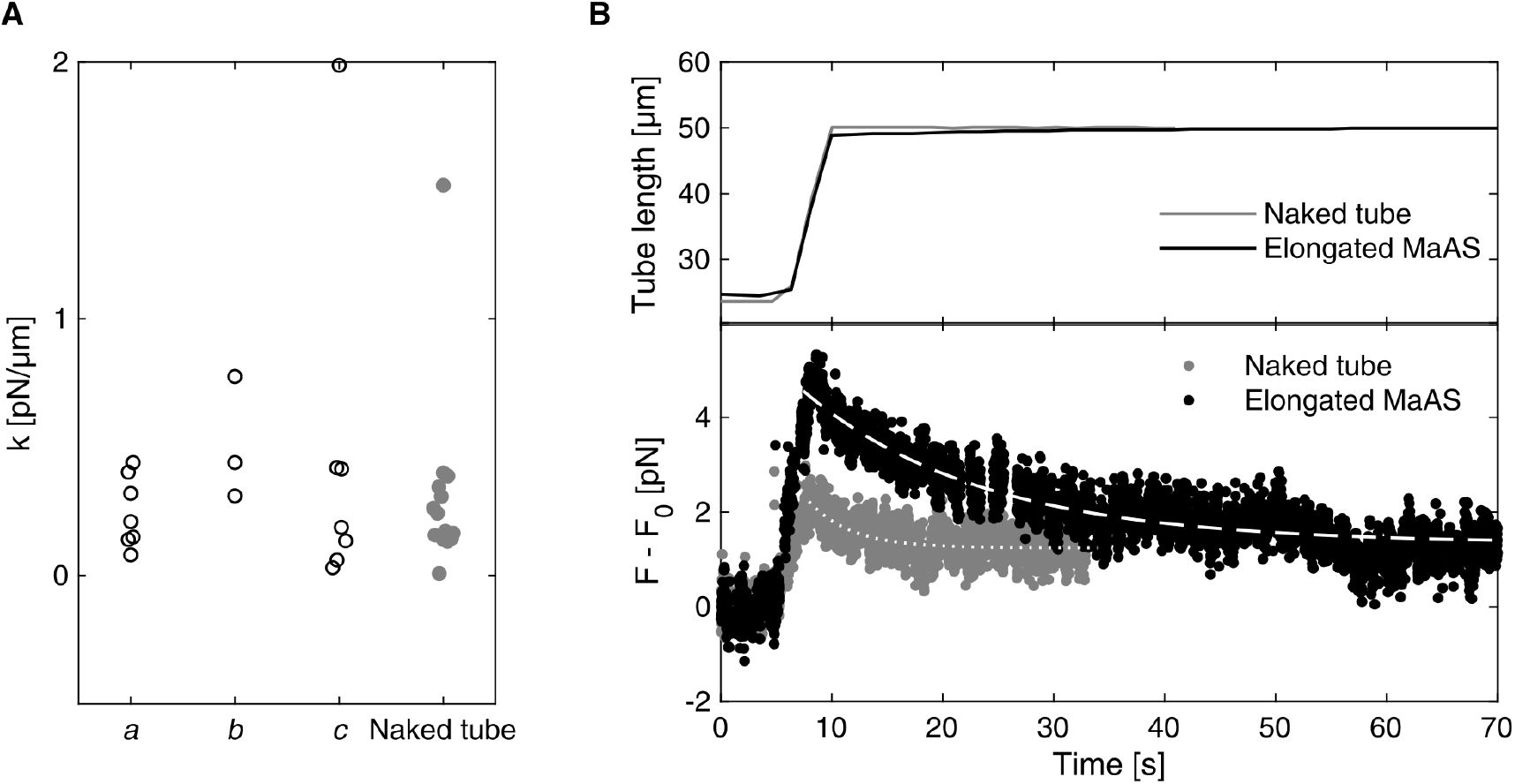
Force on an elongated MaAS. (A) Proportionality factor *k* extracted from force-elongation curves, depending on MaAS fate clarified in Fig. 3B, and for naked tubes. (B) Example of a *b* case: (top) tube length evolution as a function of time; (bottom) relative force detected from the trap, for a same tube, at different step of the injection process described in Fig. 1A: light grey corresponds to a naked tube and dark to a MaAS. Fitting curves are used to extract relaxation time (*τ*_rel_ in supplementary materials). Pushing after pulling on a naked tube is reversible.

**Fig. S5.**
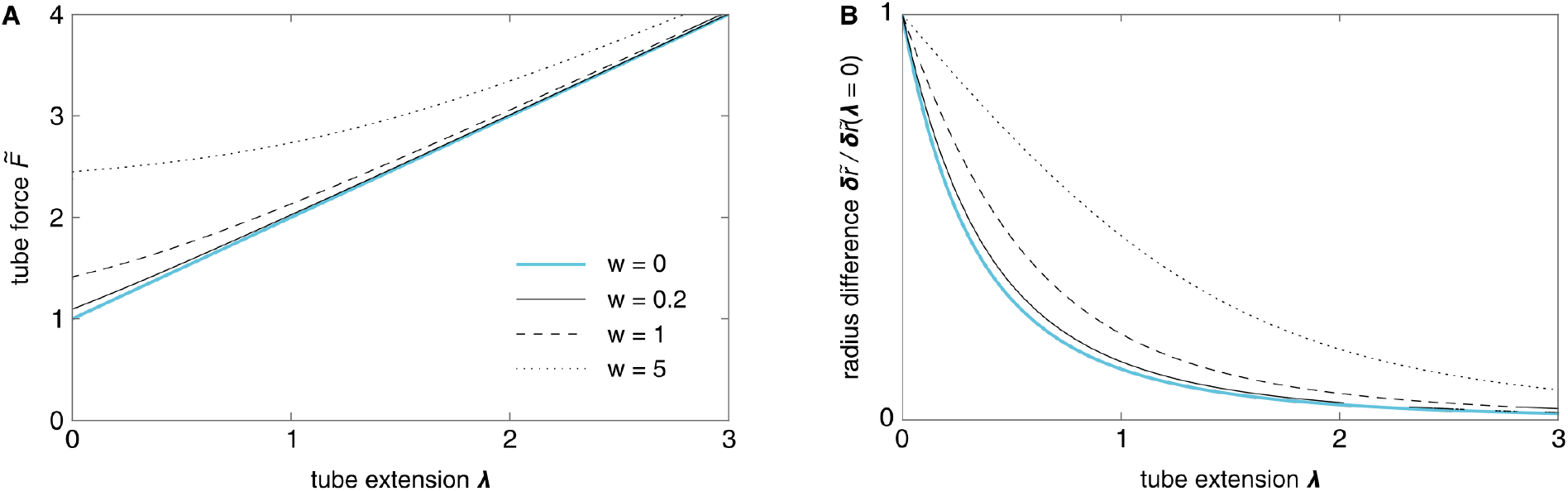
MaAS radii and force during fast pulling. Comparison of the small-*w* expressions for (A) the dimensionless force 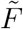 and (B) the dimensionless radius difference 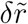 given in equation (21) (*blue lines*) with the exact expressions computed from equation (19) (*black lines*). While some quantitative discrepancies are apparent for the larger values of *w*, we find that our approximate expressions are qualitatively correct in all cases.

**Fig. S6.**
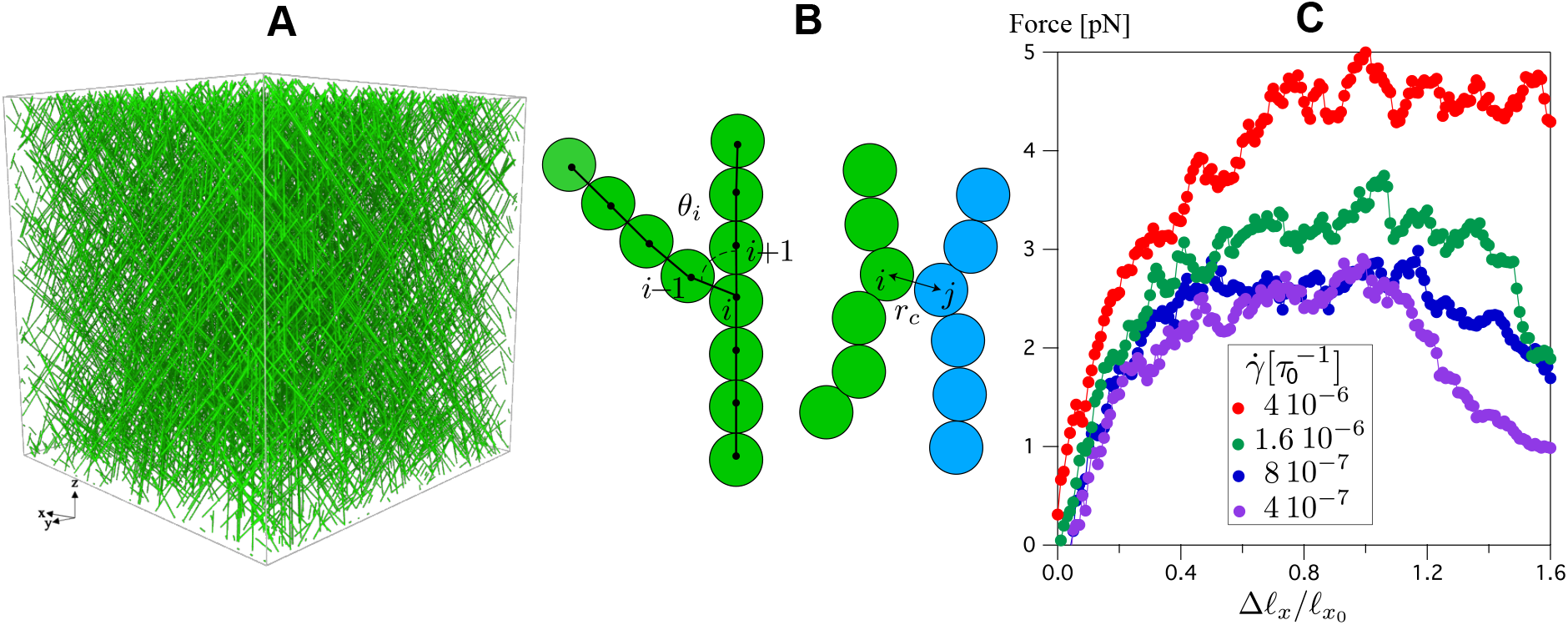
Coarse-grained simulations of branched actin. (A) Snapshot of a branched network at a constant density showing the actin filaments after the growth process along *z* axis. The branching probability is *β* = 0.03 which correspond to a network meshsize 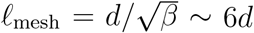. (B) Schematic representation of the discretization of the filaments into monomers of diameter *d*, the left sketch shows a branched filament. Each triplet of monomer define an angle *θ*_*i*_. The right sketch shows the exclude volume repulsive interaction between filaments with *r*_*c*_ = 2^1/6^*d* the cut-off distance. (C) Force *F* = *σ_xx_ × S* vs. deformation 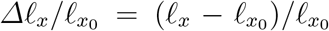 for a network of thickness *ℓ*_*z*_ = 5.5*ℓ*_mesh_. The different curves correspond to different rate of deformation 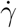.

